# Exposure to perfluorooctanoic acid accelerates *Drosophila melanogaster* juvenile development and disrupts mitochondrial metabolism

**DOI:** 10.64898/2026.06.14.730922

**Authors:** Eric A. Kilbourn, Matthew R. Lowe, Kanishka Panda, Akshay M. Bhaskaran, Guomao Zheng, Aditi R. Aalati, Adam Malave, Sophia White, Anna Graber, Nika Zulkowski, Robert Pepin, Amina Salamova, Travis Nemkov, Angelo D’Alessandro, Swathi Yadlapalli, Pramod Reddy, Edgar Meyhofer, Jason M. Tennessen

## Abstract

Per- and polyfluoroalkyl substances (PFAS) are persistent environmental contaminants with poorly understood sublethal effects on insects. Perfluorooctanoic acid (PFOA), one of the most widely distributed legacy PFAS is increasingly recognized for altering organismal physiology beyond traditional toxicity endpoints. Here, we use the fruit fly *Drosophila melanogaster* as a model to examine how PFOA exposure during larval (juvenile) development reshapes insect life-history progression and metabolic homeostasis. Our studies reveal that at environmentally relevant concentrations (nM to low µM), PFOA induces precocious expression of developmentally-regulated genes and leads to metabolic changes that persist into adulthood. At higher concentrations used to probe mechanism, PFOA accelerates larval development, disrupts mitochondrial membrane potential, and increases whole-organism metabolic heat production – results that suggest altered mitochondrial energetic efficiency. Consistent with this tradeoff, PFOA-exposed larvae that develop faster under permissive conditions exhibit heightened sensitivity to environmental stressors, including elevated temperature and reduced food hydration. Together, these findings demonstrate that PFOA disrupts metabolic and developmental processes in a dose- and context-dependent manner, highlighting sublethal effects that may influence insect resilience under environmental stress.

**SYNOPSIS STATEMENT:** Here we describe how PFOA alters the growth, development, and metabolism of the fruit fly *Drosophila melanogaster*. Specifically, we find that PFOA accelerates *Drosophila* juvenile growth while also rendering exposed larvae sensitive to environmental stress. These observations suggest that widespread PFOA contamination may impair the developmental fitness of insect populations.

## INTRODUCTION

Per- and polyfluoroalkyl substances (PFAS) are a class of synthetic chemicals widely used in industrial and consumer products for their resistance to heat, water, and chemical degradation (Buck et al., 2011). This exceptional stability has earned them the label “forever chemicals” and has led to their pervasive accumulation in the environment, with several members of this family detected across diverse ecosystems and in a wide range of organisms (Sunderland et al., 2019). In fact, the bioaccumulation of PFAS molecules is so prevalent that several individual compounds, including perfluorooctanoic acid (PFOA), are detectable in the blood of over 90% of tested individuals (Kato et al., 2011, Lewis et al., 2015, Tian et al., 2018). This is concerning given that PFAS exposure has been associated with a wide range of human health outcomes, including kidney and testicular cancers, metabolic disease, and accelerated aging (Steenland and Winquist, 2021, Fenton et al., 2021, Wee and Aris, 2023, Qi et al., 2020). Despite these concerns, the molecular and physiological mechanisms by which PFAS exposure perturbs biological systems remain incompletely understood.

While PFAS are clearly associated with human disease risk, their environmental persistence and widespread use also raise growing concerns for ecosystem health and agricultural sustainability. These concerns are particularly acute for insects, which play central roles in pollination, nutrient cycling, and food webs. Accordingly, increasing attention has turned toward understanding how PFAS contamination affects insect physiology and development. A variety of insect species have been reported to bioaccumulate PFAS through contact with contaminated soil, water, and plant surfaces (Koch et al., 2021, Judy et al., 2022, Lescord et al., 2015, Lesch et al., 2017, Lan et al., 2020, Koch et al., 2020). Moreover, PFAS exposure can disrupt key physiological processes throughout the insect life cycle – for example, PFAS have been shown to impair honeybee larval development, reduce survival, and disrupt detoxification and immune-related gene expression (Lei et al., 2025, Sonter et al., 2026). Similarly, PFAS exposure induces delayed metamorphosis in silkworms, locust, and the damselfly *Enallagma cyathigerum* (Bots et al., 2010, Wan et al., 2025), while also inducing accelerated development and maturation of the beet armyworm *Spodoptera exigua* (Omagamre et al., 2020). Together, these findings demonstrate how PFAS contaminants can disrupt insect life history events and raise the possibility that widespread PFAS pollution contributes to global insect decline.

The fruit fly *Drosophila melanogaster* has long served as a powerful genetic model for dissecting the molecular mechanisms by which toxicant exposure perturbs metabolic, physiological, and developmental processes (Rand et al., 2023). Although such studies often emphasize conserved responses that are relevant to human health (Hayot et al., 2025, Colbourne et al., 2022, Consortium, 2023), studies in the fly can also directly inform how environmental toxicants disrupt insect physiology. In particular, larval development offers a uniquely tractable framework for examining influences on the health of holometabolous insects, as the growth, development, and onset of maturation during this life stage are controlled by relatively well-defined endocrine signals and metabolic cues (Fleck et al., 2025, Dunham and Bland, 2026, Texada et al., 2020), thus enabling rapid mechanistic studies.

Here, we leverage the predictability of *Drosophila* development to demonstrate that exposure to perfluorooctanoic acid (PFOA) disrupts metabolic homeostasis and alters the rate of juvenile growth. Previous studies of *Drosophila* development have reported that PFOA exposure induces developmental delays and increases larval lethality (Luo et al., 2025, Wang et al., 2010, Yang et al., 2026, Kim et al., 2021). These findings, however, were based on exposure to relatively high PFOA concentrations, and thus left unresolved the question of how environmentally relevant doses influence *Drosophila* growth, metabolism, and developmental progression. To address this issue, we employed a two-tiered experimental framework that examine transcriptional and metabolic responses at environmentally relevant concentrations while using higher concentrations to probe underlying physiological mechanisms.

By focusing our initial analysis on ecologically relevant exposure levels, we unexpectedly observed precocious expression of late L3 gene expression programs in PFOA-exposed larvae. Subsequent dose–response analyses demonstrated that PFOA accelerates larval growth and developmental progression. Together, these findings suggest that the shift in L3 gene expression programs likely reflects accelerated developmental progression rather than direct transcriptional activation by PFOA. Moreover, this growth phenotype is highly sensitive to environmental context – when larvae are reared at elevated temperatures or on food with reduced water content, PFOA exposure instead produces the previously reported lethal phenotype, indicating that PFOA-induced developmental acceleration carries a measurable fitness cost.

Beyond its effects on developmental timing, we also find that PFOA exposure disrupts larval metabolism. PFOA-fed larvae exhibit reduced mitochondrial membrane potential and increased metabolic heat production. Strikingly, these developmental perturbations influence the adult metabolic state, as PFOA exposure during larval growth induce significant changes in the adult metabolome. Together, these findings demonstrate that developmental exposure to PFOA disrupts both life-history progression and metabolic homeostasis in *Drosophila* and underscore the need to further explore how PFAS compounds may compromise insect physiology across the life cycle.

## RESULTS

### PFOA exposure induces precocious expression of developmentally regulated genes

To assess the effects of PFOA on developing larvae, we performed RNA sequencing (RNA-seq) on L3 larvae 84 hours after egg laying (AEL) exposed to 15 ppb (0.036 µM; low-dose) and 1500 ppb (3.6 µM PFOA; high-dose) at 25 ^°^C – a timepoint near the end of the growth phase when larvae are still actively feeding (Table S1-2). Both exposure conditions induced widespread transcriptional changes, with larvae exposed to 0.036 µM or 3.6 µM PFOA exhibiting 608 and 285 differentially expressed genes (DEGs), respectively, compared with unexposed controls (adjusted *p* ≤ 0.05; log₂ fold change ≤ –1 or ≥ 1; Table S3–S4). Notably, the seemingly inverted dose-dependent relationship suggested that many of the observed transcriptional changes may reflect indirect consequences of an altered physiological or developmental state rather than a simple linear response to PFOA concentration.

Consistent with the possibility that PFOA exposure induces to a fundamental change in the progression of larval development, a closer examination of the RNA-seq data revealed that the most highly and significantly upregulated genes belonged to the salivary gland secretion (sgs) gene family (Figure 1A–B; Beckendorf and Kafatos, 1976). Members of this “glue gene” family are only expressed during a short window of larval development, immediately prior to metamorphosis, and of the top 25 DEGs, six were members of the glue gene family (*sgs1*, *sgs3*, *sgs4*, *sgs5*, *sgs7*, and *sgs8*). We independently validated these findings using a previously characterized *sgs3-GFP* reporter, which recapitulates the temporal expression pattern of the *sgs3* gene (Biyasheva et al., 2001). Consistent with the RNA-seq studies, *sgs-GFP* was robustly induced in the salivary glands of PFOA-exposed larvae at 84 h AEL but was absent in age-matched controls (Figure 1D–E).

**Figure 1.**
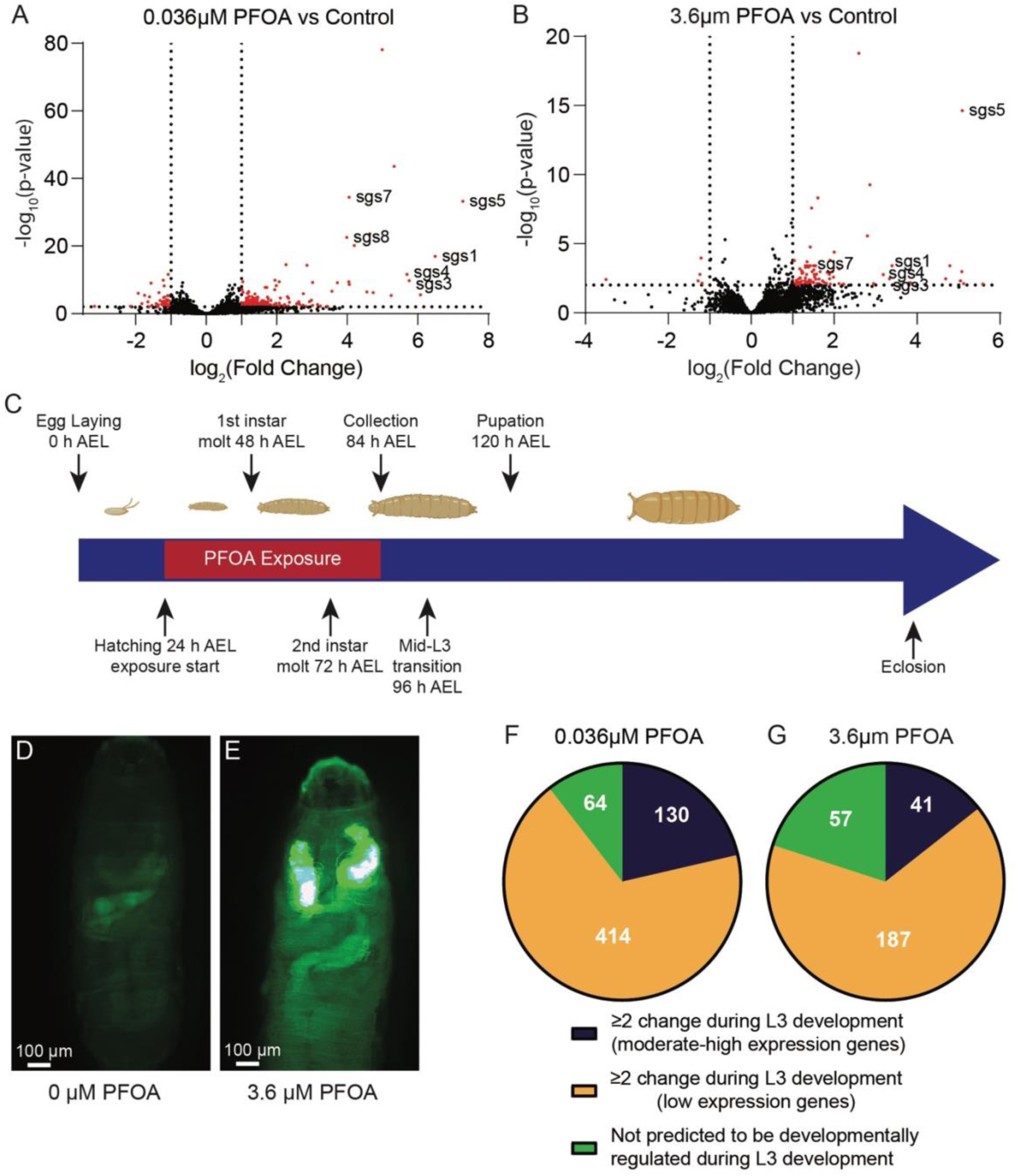
PFOA exposure induces precocious expression of developmentally regulated genes. Oregon R larvae were reared on Bloomington Drosophila Stock Center (BDSC) food containing PFOA and collected at 84 h after egg laying (AEL) for whole-body RNA-sequencing. (A–B) Volcano plots showing changes in gene expression (log₂ fold change, x-axis) and adjusted *p*-value (–log₁₀, y-axis) in larvae exposed to (A) 0.036 µM or (B) 3.6 µM PFOA relative to controls. *salivary-gland secretion* (*sgs*) genes are highlighted. (C) Schematic timeline of *Drosophila melanogaster* development indicating the PFOA exposure window, RNA-seq collection at 84 h AEL, and major developmental transitions. (D–E) Representative images of *sgs3-GFP* expression in larvae at 84 h AEL exposed to (D) 0 µM or (E) 3.6 µM PFOA. (F–G) Proportion of differentially expressed genes (DEGs) following exposure to (F) 0.036 µM or (G) 3.6 µM PFOA that correspond to genes developmentally-regulated during late L3 based on the modENCODE developmental transcriptome. DEGs are divided into three groups – not developmentally-regulated, developmentally-regulated (high expression), developmentally regulated (low expression).

Glue gene expression is a hallmark feature of a developmentally regulated event known as the mid-L3 transition (Andres et al., 1993; Biyasheva et al., 2001). Because the onset of the mid-L3 transition normally occurs shortly after the 84 h AEL collection timepoint used in this study (Figure 1C), the premature induction of glue genes in PFOA-exposed larvae suggests early activation of this developmental program. We tested this hypothesis by comparing DEGs from both exposure conditions to gene expression changes known to occur during mid-to-late L3 development (Graveley et al., 2011). Our analysis revealed that more than 75% of DEGs identified in PFOA-exposed larvae correspond to genes that undergo a ≥2-fold change following the 84 h AEL timepoint analyzed in our study (Figure 1F–G). These results suggest that the transcriptional differences observed at 84 h AEL are likely driven by accelerated developmental progression in PFOA-exposed larvae, rather than PFOA directly interfering with larval gene expression networks.

To determine whether this transcriptomic signature reflects a true acceleration of larval development, we examined the time at which metamorphosis was initiated across a range of PFOA concentrations. Our initial studies failed to detect precocious pupation at PFOA concentrations up to 100 μM (Figure 2A); however, this result was not unexpected as the timing of pupariation naturally varies among individual larvae within the same vial (McPherson et al., 2024, Andres and Thummel, 1994), and modest shifts in the onset of metamorphosis are difficult to detect. To more clearly resolve whether PFOA can influence developmental timing, we performed a dose-response analysis using higher PFOA concentrations.

**Figure 2:**
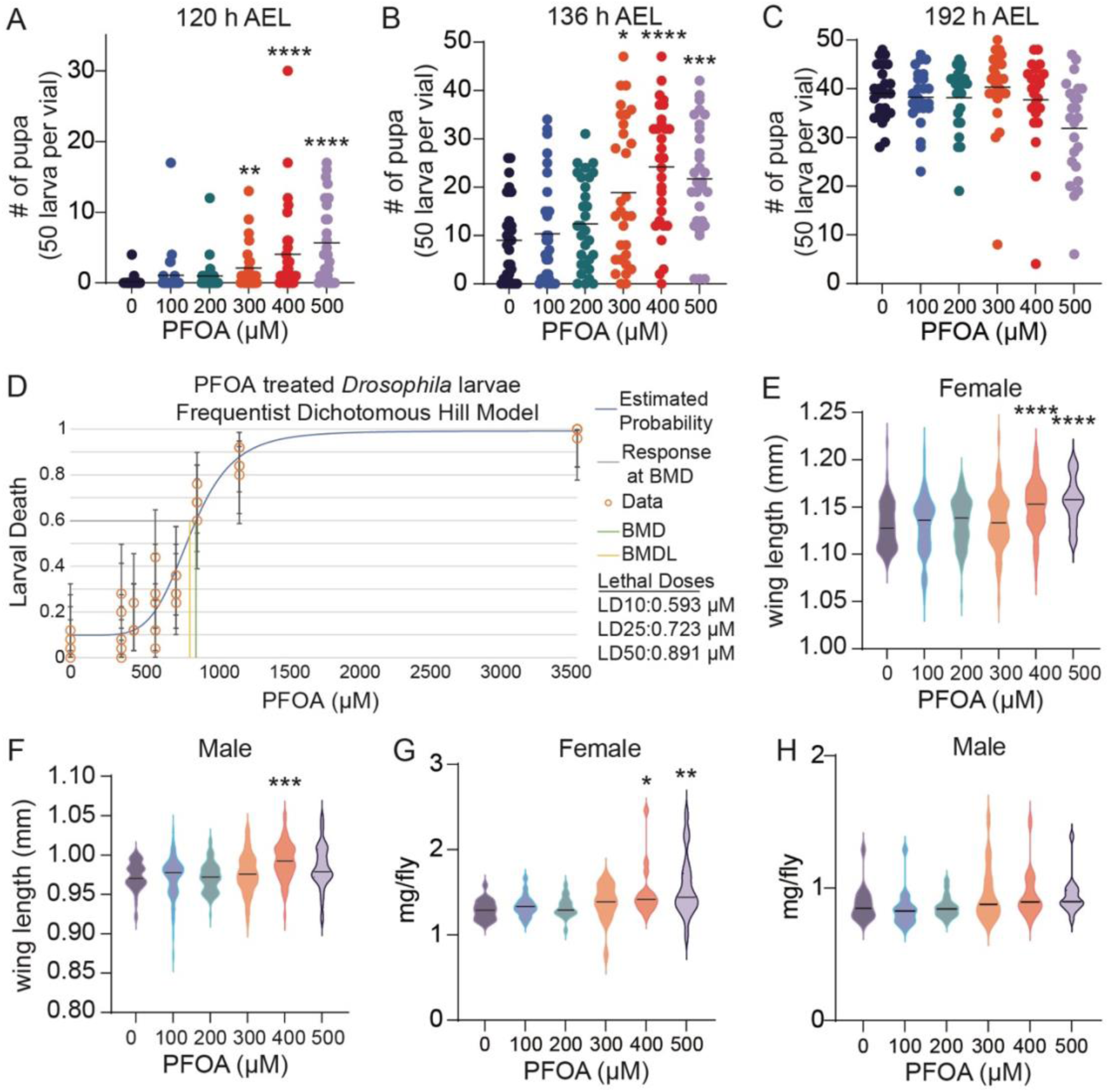
PFOA exposure accelerates larval development without reducing growth. Synchronized Oregon R embryos were placed on PFOA-containing BDSC food and the resulting larvae were raised at 25°C. (A–C) Number of pupae, including white pre-pupae, observed at (A) 120 h, (B) 136 h, and (C) 192 h after egg laying (AEL). Each data point represents an independent vial containing 50 embryos from a synchronized egg lay. (D) The estimated proportions of dead larvae at each PFOA concentration was determined based on the Dichotomous Hill model. n=4 or 5 vials with 20 flies per vial. Larval death was calculated based on the number of animals that failed to pupariate in each vial. The estimated concentration at which 10%, 25%, and 50% of each sex of flies died is indicated to the right of the graph. Lethal dose (LD), Benchmark dose (BMD), Benchmark dose low (BMDL). (E–F) Wing length of adult females (E) and males (F) measured 1 day after eclosion. Each violin plot represents measurements from 49–70 individual flies per condition. (G-H) Body mass of adult (G) females and (H) males measured 1 day after eclosion. n≤15 biological replicates per condition. Following evaluation of the data for normality assumptions, the 120 h AEL pupation data were analyzed with a Kruskal-Wallis followed by Dunn’s multiple comparisons test, while the 136 h and 192 h AEL pupation data were analyzed using a one-way ANOVA followed by Dunnett’s multiple comparisons test. Wing length measurements and fly masses were analyzed with Kruskal-Wallis followed by Dunn’s multiple comparisons test. All significance values relative to unexposed controls. \**p* < 0.05. \*\**p* < 0.01. \*\*\**p* < 0.001. \*\*\*\**p* < 0.0001.

Consistent with our RNA-seq findings, larvae reared on BDSC food containing increasing concentrations of PFOA exhibited a dose-dependent early pupariation phenotype (Figure 2A–C). At 120 h AEL, larvae exposed to 300–500 µM PFOA showed a significant increase in pupation, whereas pupariation was rarely observed in unexposed controls (Figure 2A). Although larvae exposed to 100 µM or 200 µM PFOA did not display a statistically significant increase in overall pupation frequency at this early timepoint, pupae were nevertheless consistently observed in multiple exposure vials at 120 h AEL, in contrast to controls (Figure 2A). Specifically, while only 2 of 30 control vials contained pupae at 120 h AEL, pupae were observed in 7 of 30 vials containing 100 µM PFOA and 10 of 30 vials containing 200 µM PFOA. This trend persisted at 136 h AEL, when pupation rates were significantly elevated in the 300, 400, and 500 µM PFOA exposure groups relative to unexposed larvae (Figure 2B). We would note, however, that by 192 h AEL, there was no significant difference in the pupation rate among any of the tested concentrations, indicating that the PFOA concentrations in our study did not impair overall viability under these growth conditions (Figure 2C). Consistent with this finding, subsequent dose–response assays of larval viability revealed that, in our studies, the EC10 for PFOA-induced lethality exceeds 500 µM, indicating that accelerated development occurs at concentrations well below those that limit viability (Figure 2D).

Precocious wandering and early pupariation can arise through at least two distinct physiological mechanisms. The timing of metamorphosis is governed by the integration of endocrine signals and metabolic cues, with larvae passing a key developmental checkpoint, known as “critical weight,” once sufficient biomass has been accumulated to ensure pupal and adult viability (Gillette et al., 2021; Texada et al., 2020; Tennessen and Thummel, 2011; Nijhout et al., 2014). After reaching this threshold, the timing of pupariation becomes environmentally plastic, such that stressful conditions can induce early wandering and pupation, typically at the cost of reduced growth and smaller adult size due to a shortened feeding period (Beadle et al., 1938; Nijhout et al., 2014; Mirth et al., 2005). We did not observe this outcome following PFOA exposure. Instead, adult body mass and wing length were unchanged at lower exposure levels and increased at higher concentrations, with females reared on 400 µM and 500 µM PFOA and males exposed to 400 µM PFOA exhibiting increased wing length relative to controls (Figure 2E–H). Together, these findings argue against stress-induced acceleration of metamorphosis and instead support a model in which PFOA exposure accelerates larval growth in a dose-responsive manner.

### Larval PFOA exposure reduces mitochondrial membrane potential and induces excess heat generation

Relatively few genetic or metabolic manipulations are known to accelerate *Drosophila* larval development. Among the best characterized is increased insulin signaling, which promotes growth and can shorten developmental time (Ikeya et al., 2002; Brogiolo et al., 2001; Dunham and Bland, 2026; Walkiewicz and Stern, 2009). Insulin pathway activity is closely linked to the subcellular localization of the transcription factor FOXO, which translocates from the cytoplasm to the nucleus in response to reduced insulin signaling (Jünger et al., 2003; Puig et al., 2003). We therefore asked whether PFOA accelerates development by enhancing insulin signaling, predicting that PFOA exposure would decrease the nuclear-to-cytosolic FOXO ratio. However, FOXO localization in PFOA-exposed larvae at 84 h AEL was indistinguishable from that observed in unexposed larvae (Figure S1A-B), indicating that PFOA does not enhance insulin/FOXO signaling axis in the larval fat body. Similarly, a reanalysis of the RNA-seq data further supported this conclusion, as no significant enrichment of insulin- or FOXO-related genes was detected among the DEGs (Table S5). Together, these findings suggest that PFOA-induced acceleration of larval development is not mediated by canonical insulin/FOXO signaling.

Because insulin signaling does not appear to account for the observed developmental acceleration, we next examined whether PFOA exposure is associated with alterations in mitochondrial metabolism, a process previously linked to larval growth rate. For example, ectopic expression of the alternative oxidase (AOX) from *Ciona intestinalis* accelerates *Drosophila* larval development and is associated with altered mitochondrial electron transport, changes in redox balance, and increased metabolic flux (Garcia et al., 2025). Together with prior findings that PFOA reduces mitochondrial membrane potential in mouse embryos (Zhou et al., 2022), these observations led us to hypothesize that PFOA exposure is associated with altered mitochondrial metabolism in *Drosophila* larvae. To test this possibility, we assessed mitochondrial membrane polarization in the larval fat body using the fluorescent dye JC-1, which accumulates within mitochondria in a membrane potential–dependent manner. Under conditions of high membrane potential, JC-1 forms red-fluorescent J-aggregates, whereas loss of membrane potential favors the monomeric, green-fluorescent form (Smiley et al., 1991). Fat bodies dissected from L3 larvae exposed to 500 µM PFOA at 25 °C displayed a significant reduction in the ratio of red (568 nm) to green (488 nm) fluorescence relative to unexposed controls, indicating a loss of mitochondrial membrane potential (Figure 3A–B).

**Figure 3.**
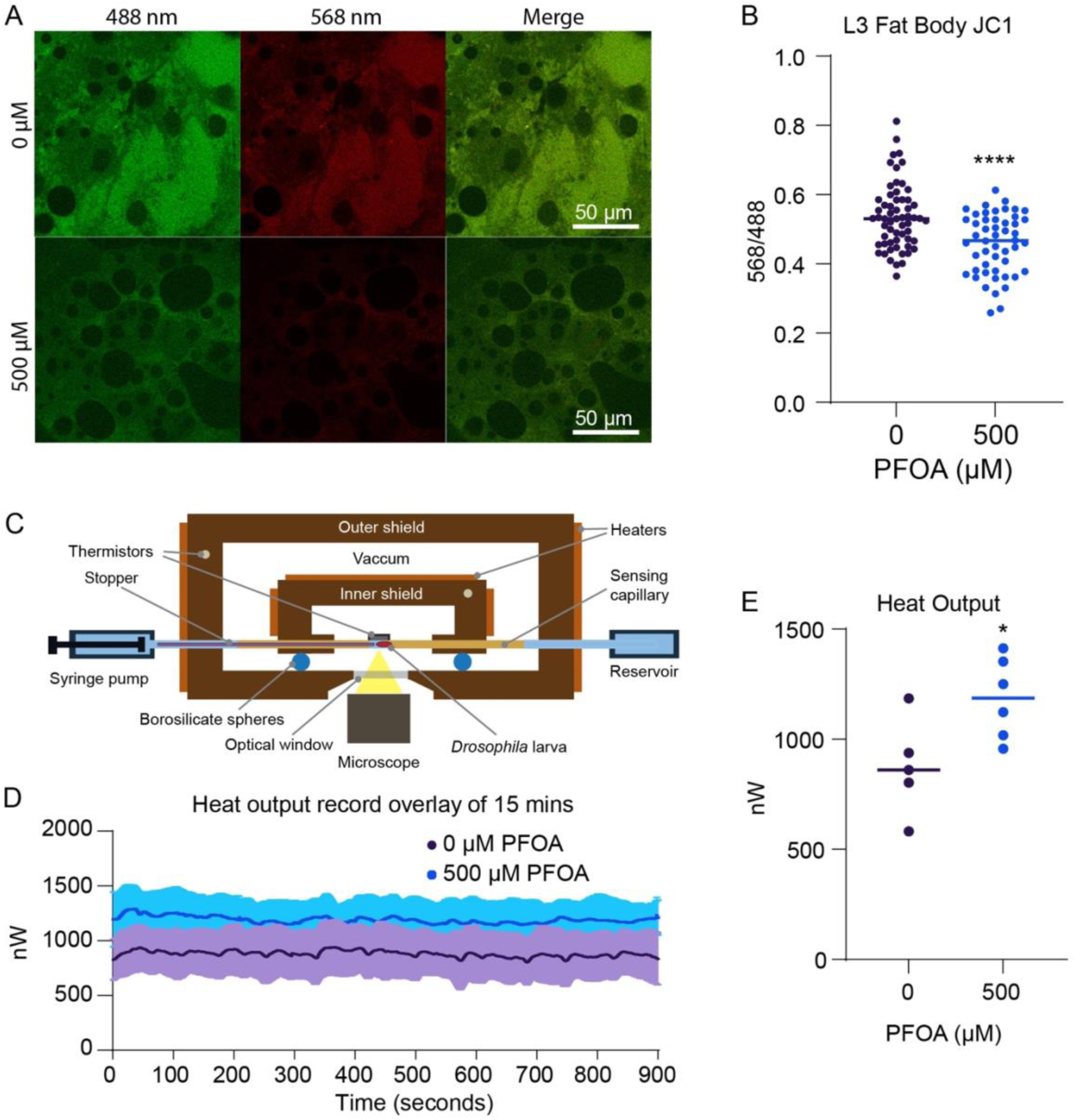
PFOA exposure alters mitochondrial membrane potential and increases heat production. Fat bodies were dissected from L3 larvae exposed to 0 or 500 µM PFOA at 84 h after egg laying (AEL), stained with JC-1, and imaged live. (A) Representative fat body images showing JC-1 fluorescence in the 488 nm (monomer), 568 nm (J-aggregate), and merged channels. (B) Quantification of the JC-1 568/488 fluorescence ratio, consistent with reduced mitochondrial membrane potential following PFOA exposure. Each data point represents a field of view from 5 fat bodies, with fluorescence ratios averaged across multiple regions of interest. Similar results were obtained from 4 independent experiments. (C) Schematic diagram of the biocalorimetry platform used to measure larval heat output. (D) Overlay of 15 minutes of stable heat output readings from representative larvae. (E) Quantification of heat output from L3 larvae (84–90 h AEL) that were resynchronized at the L2–L3 molt. Each data point represents an individual larva. For (B) and (E), data were analyzed using unpaired Student’s t-tests after assessing normality and variance. \**p* < 0.05, \*\*\*\**p* < 0.0001.

To determine whether disruption of mitochondrial membrane potential was associated with altered energetic output, we measured metabolic heat production of the larvae using a high-resolution biocalorimetry platform (Figure 3C). This platform has been described and used in past work to study metabolism in *Drosophila* tissues (Panda et al., 2025). Using this approach, we found that L3 larvae synchronized at the L2/L3 molt and exposed to 500 µM PFOA produced ∼37% more heat than unexposed controls (Figure 3D–E), consistent with increased metabolic flux. This increase in heat output corresponded to a modest absolute temperature change of approximately 1 mK, which is unlikely to directly accelerate development through thermal mechanisms. Instead, these data support a model in which PFOA exposure reduces mitochondrial energetic efficiency, accompanied by a compensatory increase in larval metabolic flux.

### PFOA sensitize larval development to environmental stress

Our findings demonstrate that PFOA exposure increases the larval metabolic rate and accelerates larval growth. While such phenotypes might seem advantageous, we hypothesized that this altered physiological state would limit the ability of developing larvae to tolerate environmental stress. We first tested this idea by examining larvae reared at 29 °C, near the upper limit of temperatures permissive for *Drosophila* development. Under these conditions, PFOA-exposed larvae still pupariated earlier than unexposed controls (Figure 4A), indicating that PFOA-induced accelerated growth still occurs at this temperature. PFOA exposure also remained associated with increased overall growth, with the wing lengths of 200-500 µM PFOA exposed females, and 300 µM PFOA exposed males (Figure S2A-B). However, at 29 °C a substantial fraction of larvae raised on PFOA-supplemented food wandered away from the food and died on the sides of the vial, with increased PFOA doses inducing higher levels of lethality (Figure 4B). Thus, higher concentrations of PFOA render larvae sensitive to heat stress.

**Figure 4.**
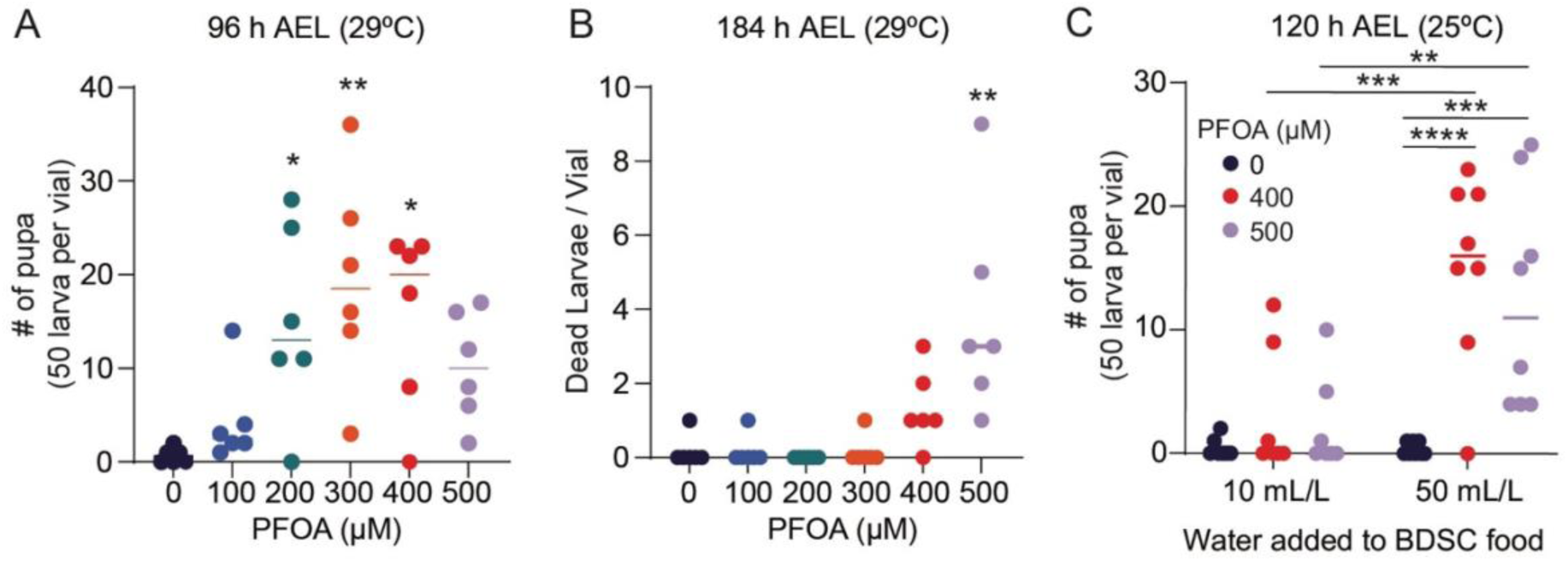
PFOA exposure sensitizes larval development to environmental stressors. (A,B) Synchronized Oregon R embryos were placed on PFOA-containing BDSC food and the resulting larvae were raised at 29 °C to assess sensitivity to environmental stress. (A) Number of pupae, including white pre-pupae, observed at 96 h AEL following larval exposure. (B) Number of small wandering larvae that died on the sides of vials 184 h AEL. Data were analyzed using a one-way ANOVA followed by Dunnett’s multiple comparisons test. Significance values relative to the unexposed control. (C) Comparison of pupation rates on summer versus winter BDSC food formulations during November and December 2025, reflecting differences in food hydration. Data were analyzed using a two-way ANOVA followed by a Tukey’s multiple comparisons test. For all panels, each data point represents a biological replicate containing 50 synchronized embryos. \**p* < 0.05. \*\**p* < 0.01. \*\*\**p* < 0.001. \*\*\*\**p* < 0.0001.

In addition to its effects on heat tolerance, we serendipitously found that PFOA also sensitizes larval development to dehydration. All media used in this study were prepared by the Bloomington Drosophila Stock Center (BDSC) prior to PFOA supplementation. The BDSC recipe is adjusted seasonally, with additional water added to the food during the winter months as a means of compensating for lower ambient humidity and increased evaporation. During the course of our studies, we noticed that PFOA-exposed larvae exhibited high levels of lethality in October and November, prior to the switch to the winter recipe. Since the seasonal BDSC formulations are otherwise identical, we tested whether food moisture accounts for this phenomenon. We tested this hypothesis in November 2025 by supplementing BDSC summer food with the additional water used in the winter recipe. This modification significantly altered PFOA-associated outcomes (Figure 4C), with increased moisture content leading to improved larval viability and accelerated developmental rates. These findings not only demonstrate that food hydration state is a critical modifier of the PFOA-induced phenotypes but also suggest that subtle differences in the larval food composition and hydration might explain the discrepancies between our results and earlier studies, which reported that PFOA induces developmental delays and lethality. Together, our results show that PFOA can induce accelerated development in optimal conditions but also sensitizes exposed larvae to environmental stress.

### PFOA exposure during larval development results in persistent metabolic reprogramming

The elevated metabolic flux observed in PFOA-exposed larvae at higher concentrations led us to ask whether environmentally relevant exposures similarly perturb larval metabolic state during development. To address this, we profiled the larval metabolome following low-dose PFOA exposure (0.036 µM and 3.6 µM), sampling larvae at 84 h AEL (Table S6). We observed relatively few metabolic changes (Figure S3A-B), suggesting that low-dose PFOA has limited immediate effects on steady-state larval metabolism. Notably, however, 2-hydroxyglutarate (2HG) was elevated with PFOA exposure (Figure S3B). Because L-2HG is the predominant enantiomer in larvae and its accumulation is strongly promoted by mitochondrial redox imbalance (Li et al., 2017, Mahmoudzadeh et al., 2024, Mahmoudzadeh et al., 2020), this pattern is consistent with subtle alterations in redox-linked metabolic processes that parallel mitochondrial changes observed at higher PFOA concentrations.

As an extension of these findings, we asked whether larval PFOA exposure produces metabolic alterations in subsequent life stages. Because PFOA accumulated during larval development may, depending on dose, persist through metamorphosis, developmental exposure could result in prolonged internal exposure later in the life cycle. To test this possibility, we reared larvae on food containing 3.6 µM, 36 µM, or 360 µM PFOA, and newly eclosed adults were maintained on untreated food for five days prior to analysis (Figure 5A). LC-MS-based measurements revealed detectable PFOA accumulation in adults derived from larvae exposed to 36 µM and 360 µM PFOA, whereas PFOA was not detectable in adults exposed to 3.6 µM during development at the timepoints analyzed (Figure 5B–C). These results indicate that larvae bioaccumulate PFOA in a dose-dependent manner and that retained PFOA can persist into adulthood following higher-dose developmental exposure. However, PFOA levels in adults derived from lower-dose exposures were below the limit of detection.

**Figure 5.**
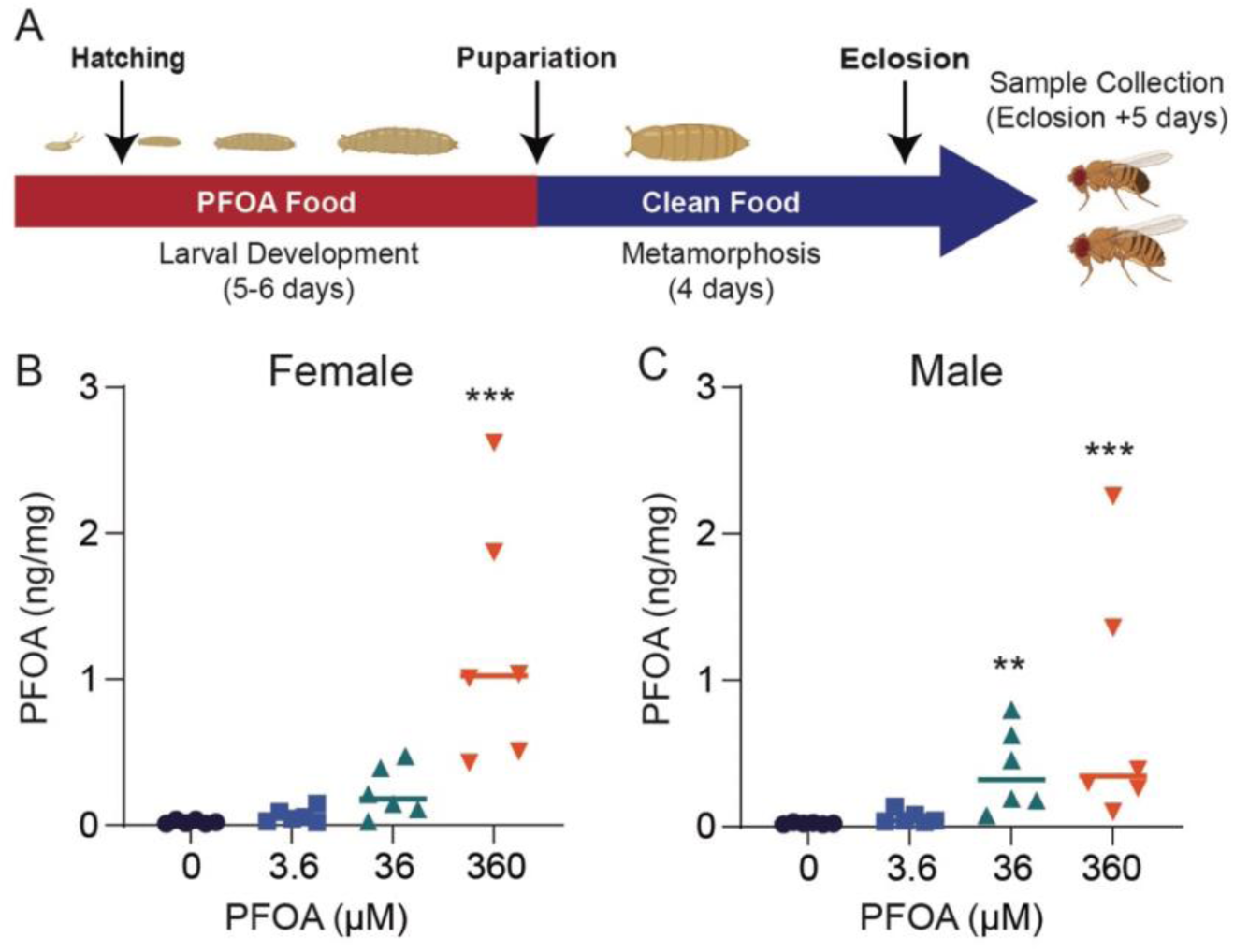
Developmental PFOA exposure leads to dose-dependent retention in adult *Drosophila*. (A) Schematic illustrating the exposure paradigm. Embryos were allowed to hatch on Bloomington Drosophila Stock Center (BDSC) food with increasing PFOA concentrations. Upon pupariation, both control and exposed animals were transferred to fresh vials of BDSC food containing no PFOA. Males and females were collected 5 days post-eclosion, sorted by sex, and PFOA concentrations were measured by UPLC–MS in (B) adult males and (C) females. Data represent n = 6 biological replicates per condition, with 3–12 flies per replicate and are normalized to sample mass. Female data were analyzed using one-way ANOVA followed by Dunnett’s multiple-comparison test, while male data were analyzed using a Kruskal–Wallis test followed by Dunn’s multiple-comparison test. \*\**p* < 0.01. \*\*\**p* < 0.001.

Using the same developmental exposure paradigm described above (Figure 5A), we performed a semi-targeted metabolomic analysis of adults collected five days post eclosion. This analysis revealed that, despite the absence of detectable PFOA following low-dose larval exposures (Figure 5B-C), both 0.036 µM and 3.6 µM larval treatments markedly altered the adult metabolome (Table S7). Partial least-squares discriminant analysis (PLS-DA) revealed clear separation between PFOA-exposed and control adults (Figure 6A–B), indicating that developmental exposure induces persistent metabolic changes. Examination of the most significantly altered metabolites in each sex revealed disruptions in carbohydrate and mitochondrial metabolism, including reduced pyruvate and acetoacetate and modest increases in ATP (Figure 6C–D). The most pronounced effects, however, were observed among acyl-carnitine species. Across sexes, 12 acyl-carnitines were among the top 20 significantly effected metabolites, five of which were elevated in both males and females (Figure 6C–D). This pattern was confirmed by targeted LC–MS analysis of AC(16:0) and AC(18:1) – two of the most abundant acyl-carnitines in *Drosophila* that have been the focus of previous fly metabolic studies (Fasteen et al., 2025, Lam et al., 2020, Palanker Musselman et al., 2016). Consistent with the semi-targeted analyses, both species were elevated following larval exposure to 3.6 µM PFOA, however, we note that the abundance of AC(16:0) and AC(18:1) in 0.036 µM exposure samples were not significantly different than the unexposed controls (Figure 7A–D). Overall, our findings demonstrate that PFOA exposure during development can induce persistent metabolic reprogramming into adulthood, with notable effects on fatty acid and mitochondrial metabolism.

**Figure 6.**
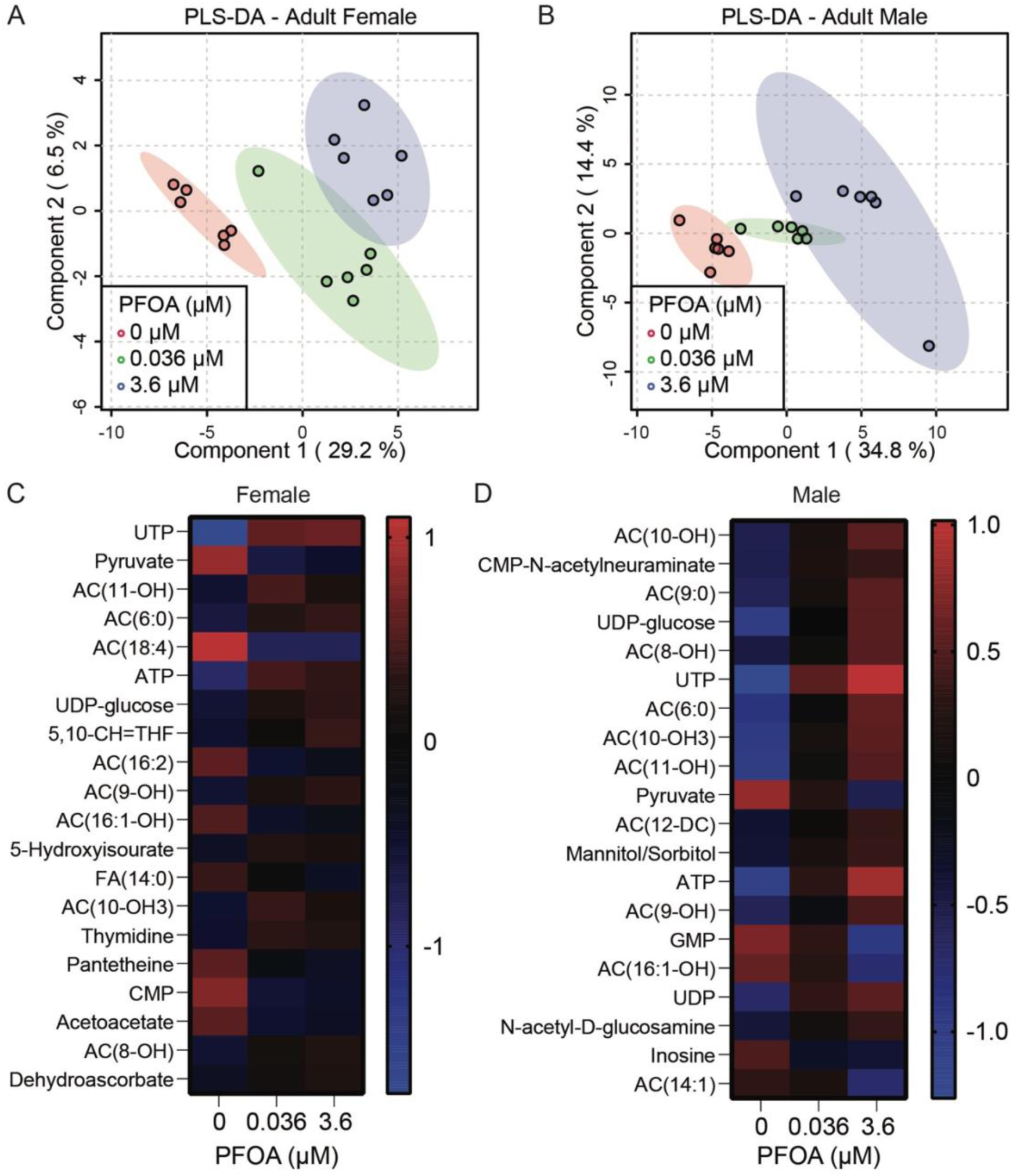
Developmental PFOA exposure induces persistent alterations in adult metabolism. Semi-targeted metabolomic analysis was performed on adults derived from exposed larvae. (A–B) Partial least squares discriminant analysis (PLS-DA) of metabolomic data from 5-day-old adult (A) females and (B) males following larval exposure to 0, 0.036, or 3.6 µM PFOA. (C–D) Heatmaps showing the most statistically significant metabolites in adult (C) females and (D) males following developmental PFOA exposure. Data were first normalized to sample mass and the spike-in internal d4-succinic acid standard followed by log normalization and Pareto scaling. Data analysis conducted with Metaboanalyst 6.0.

**Figure 7.**
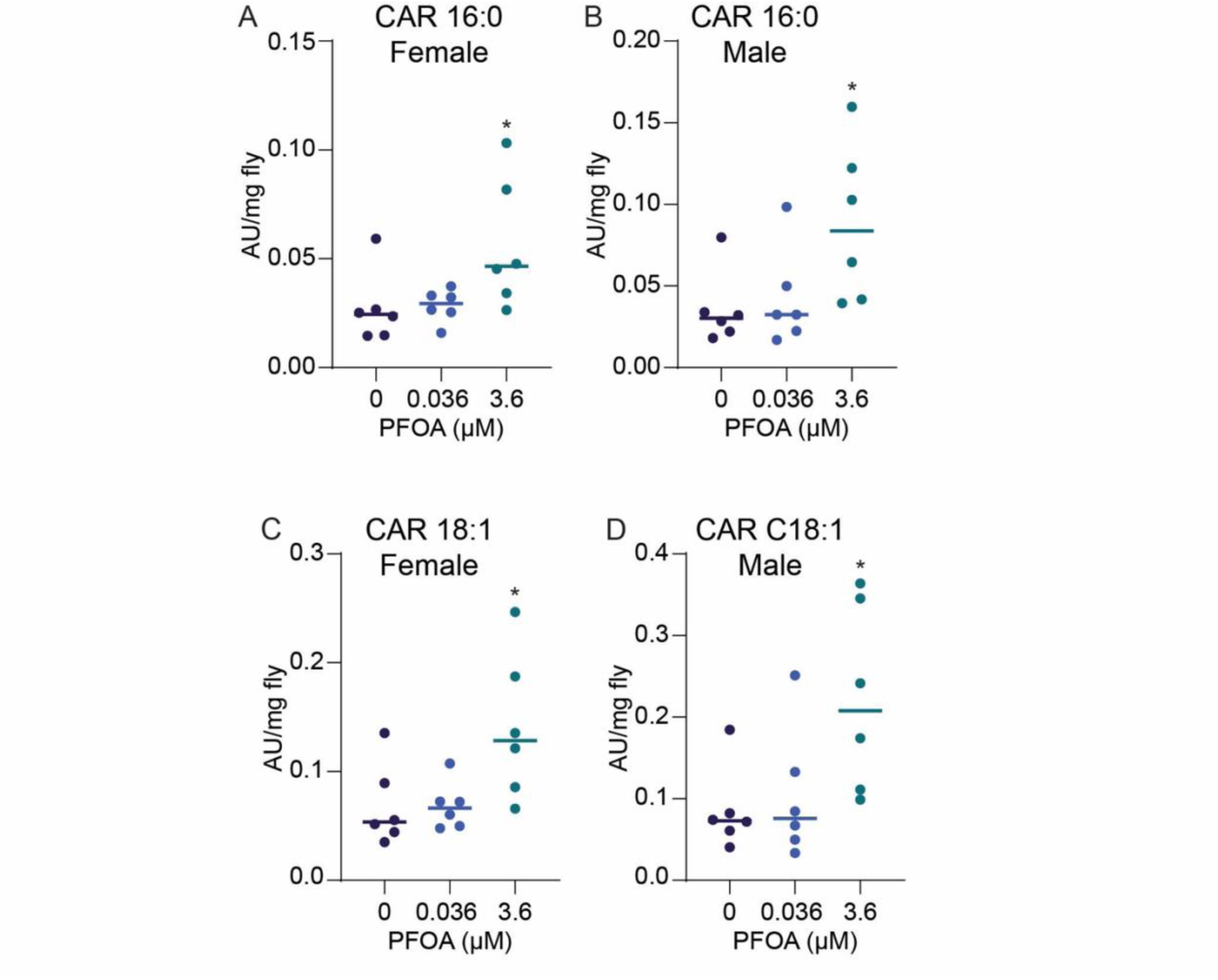
PFOA larval exposure induces elevated acyl-carnitine levels in adult flies. Larvae were exposed to 0, 0.036, or 3.6 µM PFOA and the resulting adult female and male flies were analyzed 5 days after eclosion using targeted LC-MS analysis to measure the relative abundance of (A-B) palmitoyl-L-carnitine (CAR 16:0) and (C-D) oleoyl-L-carnitine (CAR 18:1). For all analyses, *n* = 6 biological replicates per condition, 20 flies per replicate. Statistical significance for targeted analyses was determined using a one-way ANOVA followed by Dunnett’s multiple comparisons test for the female CAR 18:1 analysis and Kruskal-Wallis test followed by a Dunn’s multiple comparisons test for the male CAR 18:1, and male and female CAR 16:0. Significance values relative to the unexposed controls. \**p* < 0.05.

## DISCUSSION

Here we demonstrate that PFOA exposure can accelerate *Drosophila melanogaster* larval development and inducing long-lasting metabolic alterations. Moreover, these phenotypes are dose-responsive, with higher concentrations inducing early pupariation, reducing mitochondrial membrane potential, increasing heat production, and heightening larval sensitivity to environmental stress. While our mechanistic studies rely on higher concentrations, the basic observation that PFOA accelerates larval development and alters fatty acid metabolism are evident at the lower, environmentally relevant concentrations. Together, these observations demonstrate that PFOA significantly interferes with *Drosophila* growth, development, and metabolism across a wide range of exposure concentrations.

At first glance, these findings appear to contradict earlier *Drosophila* studies that reported developmental delays and increased lethality following PFAS exposure (Luo et al., 2025, Wang et al., 2010, Yang et al., 2026, Kim et al., 2021). However, this apparent contradiction can be explained by the influence of environmental variables on the PFOA phenotype, as we found that modest changes in rearing conditions, such as food hydration and temperature, can shift outcomes from accelerated development to impaired growth and lethality. Consistent with this idea, even small increases in environmental temperature have been shown to markedly exacerbate the physiological and fitness costs of otherwise sublethal chemical exposures during *Drosophila* larval development (Gandara et al., 2024).

Our transcriptomic analysis further supports a model in which PFOA alters developmental timing rather than directly regulating gene expression. Because samples were collected at a fixed time point, the observed precocious expression of stage-specific gene expression programs is best explained by advancement in developmental stage rather than widespread transcriptional activation by PFOA. Moreover, similar outcomes have been reported in other insect systems, including accelerated development of the beet armyworm *Spodoptera exigua* following exposure to the PFAS perfluorobutanoic acid (Omagamre et al., 2020). Together, these observations suggest that PFAS compounds can, under permissive conditions, promote accelerated insect development but also render the exposed animals sensitive to environmental stress.

Multiple signaling pathways are known to coordinate larval growth rate with developmental timing, most prominently insulin signaling (Fleck et al., 2025, Gillette et al., 2021, Texada et al., 2020). However, our data do not support a major role for canonical insulin/FOXO signaling in mediating the PFOA phenotype. Nuclear FOXO localization in the larval fat body was not reduced in PFOA-exposed animals, and transcriptomic analyses revealed little overlap between PFOA-responsive genes and curated insulin pathway gene sets. While these results do not exclude subtle, tissue-specific, or transient alterations in insulin signaling, they argue against hyperactivation of this pathway as the primary driver of accelerated development.

Instead of altered insulin signaling, our findings point to altered mitochondrial function and elevated metabolic rate as key features of the larval response to PFOA. At higher concentrations, PFOA exposure reduced mitochondrial membrane potential in the larval fat body while simultaneously increasing whole-animal heat production, consistent with disrupted mitochondrial metabolism accompanied by compensatory increases in metabolic flux. In parallel, environmentally relevant exposures altered acyl-carnitine abundance, suggesting perturbations in mitochondrial lipid handling and oxidative metabolism. Together, these developmental and metabolic phenotypes parallel those observed in larvae in which mitochondrial redox balance is genetically perturbed. For example, expression of the alternative oxidase (AOX), which provides an alternative route for electron flow through the mitochondrial electron transport chain and alters energetic efficiency, accelerates larval development and increases metabolic rate (Garcia et al., 2025). Similarly, knockdown of the mitochondrial transcription factor A homolog (TFAM), which reduces mitochondrial DNA expression and electron transport chain capacity, has been shown to induce glycolytic enzyme expression and accelerate larval development (Sriskanthadevan-Pirahas et al., 2025, Sriskanthadevan-Pirahas et al., 2022). Overexpression of lactate dehydrogenase (Ldh) in the larval fat body also accelerates development (Sriskanthadevan-Pirahas et al., 2022), supporting the idea that larval growth rate can be influenced by changes in cellular redox balance. Thus, while PFOA clearly disrupts mitochondrial metabolism, we propose that altered mitochondrial function actively contributes to broader metabolic reprogramming that accelerates growth while reducing physiological flexibility and stress tolerance.

Beyond its effects on larval development, a key finding of this study is that PFOA exposure during development produces persistent alterations in adult metabolism. While low-dose PFOA exposure induced relatively modest metabolic changes in larvae, adults that developed from exposed larvae exhibited a substantially larger number of metabolic alterations, including elevated acyl-carnitine levels consistent with disrupted mitochondrial fatty acid oxidation. Importantly, these adult metabolic phenotypes were observed even in the absence of detectable PFOA in adults following low-dose larval exposure. There are at least two, not mutually exclusive, explanations for this observation. First, it is possible that low concentrations of PFOA acquired during larval development persist into adulthood at levels below the limit of detection, and that this residual internal exposure is sufficient to drive the observed metabolic differences. Under this model, adult phenotypes reflect the cumulative effects of prolonged, low-level internal exposure initiated during development. Alternatively, these findings may reflect true metabolic reprogramming during larval development, whereby transient PFOA exposure permanently alters developmental trajectories or metabolic setpoints that persist into adulthood, even after the chemical itself has been eliminated. Distinguishing between these possibilities will be an important goal for future studies.

More broadly, our findings suggest that chronic, low-level PFAS contamination may influence insect populations in ways that are not readily captured by traditional toxicity endpoints. Rather than acting exclusively as classical toxicants that slow development or increase acute mortality, our studies in *Drosophila* reveal that PFOA is capable of accelerating growth under permissive conditions while reducing resilience to environmental stress, raising the possibility that similar life-history tradeoffs could occur in other insect species. Given the central roles insects play in ecosystem function, such effects could contribute to population-level vulnerability over time. Together, our observations in the fly underscore how chronic PFOA exposure may reconfigure insect life-history strategies in ways that disrupt the developmental plasticity required to persist under increasingly variable environmental conditions.

Finally, this work establishes *Drosophila melanogaster* as a powerful genetic model system for dissecting mechanisms of PFAS toxicity. While this organism does not capture the full ecological complexity experienced by insects in natural environments, the suite of genetic tools available to study toxicological responses in this fly is unparalleled and enables experimental approaches that can guide hypothesis-driven studies in other insect systems. As PFAS contamination remains widespread and persistent, such mechanistic approaches will be essential for understanding how these ‘forever chemicals’ influence development, metabolism, and fitness.

## METHODS

### Drosophila husbandry and genetics

Fly stocks were maintained on standard Bloomington Drosophila Stock Center (BDSC) food as described (Holsopple et al., 2023). Unless otherwise noted, cultures were maintained at 25 °C with 50% relative humidity under a 12 h light/12 h dark cycle. The following BDSC stocks were used in this study: Oregon-R (RRID:BDSC_25211), and *w[*]; P{w[+mC]=Sgs3-GFP}3* (RRID:BDSC_5885). Flybase was used as a reference throughout the study (Öztürk-Çolak et al., 2024).

### PFOA food preparation

For experiments involving PFOA concentrations ≤ 3.6 µM, a 1.5 g L⁻¹ stock solution was prepared by dissolving powdered PFOA (MilliPore Sigma, 171468-5G) in Milli-Q water. For higher PFOA concentrations, solid PFOA was dissolved directly into liquid BDSC food, with Milli-Q water added to a final volume equal to 1% of the total food volume. Prepared food was dispensed into either vials (10 mL per vial; Genessee, 32-116), 33-mm Petri dishes (10 mL/dish; CELLTREAT, 229638) or 6oz bottles (50 mL per bottle; Genessee, 32-130). Food was stored covered at 4 °C until use.

### Preparation of exposure chambers

Exposure chambers were prepared using 55-mm Petri dishes (Fisher Scientific, FB012921). Holes were poked in the lids using a heated dissecting needle (Fisher Scientific, 13-820-024), and a fine mesh was secured over the openings on the interior side of each lid using nail polish. A circular piece of Whatman No. 1 filter paper (WHA1001055) was placed in the bottom of each Petri dish and moistened with 0.5 mL of 1x PBS. Half of a 33-mm Petri dish of food was added to each chamber. After embryos or larvae were transferred to the food, chambers were sealed with Parafilm prior to placing in an incubator.

### PFOA embryo and larval exposures, developmental timing assays, and adult phenotyping

Mating bottles of adult flies that were maintained in a 12 h light/12 h dark cycle were subjected to a 2 h pre-lay period to prevent egg retention, followed by a 4 h egg-lay from which embryos were collected. For embryonic exposures, embryos were transferred directly onto PFOA-supplemented BDSC food. For first-instar (L1) exposures, embryos were maintained for 24 h on molasses–yeast egg-laying caps in Petri dishes before transfer to PFOA-spiked food within 4 h of hatching. Unless otherwise indicated, exposures were conducted in custom exposure chambers (described above) containing BDSC food supplemented with the indicated PFOA concentration.

Larval lethality was determined by exposing replicates of 50 embryos in exposure chambers to varying concentrations of PFOA in Bloomington food and counting the number of larvae that made it to pupation. BMD and lethal dose curves were generated using the EPA BMD3 software (Agency, 2023). The parameters used were dichotomous analysis, dichotomous hill and added risk for BMDs of 0.1, 0.25 and 0.5.

Developmental timing assays were conducted by placing synchronized groups of 50 embryos on PFOA-supplemented food and the number of white pre-pupa and pupa were scored at 120 h, 136 h, and 192 h AEL.

For adult phenotypic measurements, newly eclosed flies transferred to untreated BDSC food, aged for 24 h, placed in pre-tared 1.5 mL microcentrifuge tubes, and flash-frozen in liquid nitrogen. Sample mass was determined by allowing the unopened frozen tubes to return to room temperature, after any remaining condensation on the outside of the tube we removed with a paper towel and the mass measured using a Mettler Toledo XS105 Analytical Balance. Similarly, wing measurements were conducted by removing a single wing from each frozen fly. The wing was subsequently mounted on a slide in 80% glycerol and imaged using a Leica MZ10F stereomicroscope equipped with an MC170 HD camera. Wing length was measured using ImageJ as the distance from the distal end of longitudinal vein L3 to the junction of veins L2 and L3.

### RNA-seq analysis

Synchronized populations of embryos were collected as described above. First-instar (L1) larvae were collected within 4 h of hatching and transferred to exposure chambers containing BDSC food supplemented with 0, 0.036, or 3.6 µM PFOA (25 larvae per chamber; three chambers per concentration). Sixty hours after hatching (84 h after egg laying [AEL]), third-instar (L3) larvae were collected from individual exposure chambers, transferred to 1.5-mL microcentrifuge tubes, washed three times with ice-cold 1× PBS (pH 7.4), and flash-frozen in liquid nitrogen. RNA was extracted by grinding frozen larval pellets in TRIzol Reagent (Thermo Fisher Scientific, 15596018) using a pellet pestle, following the manufacturer’s instructions. Purified RNA was resuspended in 50 µL of nuclease-free water (HyPure, SH30538.02) and subsequently repurified using a Qiagen RNeasy Plus Micro Kit. Sequencing and data analysis was conducted in the Indiana University Center for Genomics and Bioinformatics.

Reads were adapter trimmed and quality filtered using Trimmomatic ver. 0.38 (Bolger et al., 2014) setting the cutoff threshold for average base quality score at 20 over a window of 3 bases, excluding the reads shorter than 20 bases post-trimming (parameters: LEADING:20 TRAILING:20 SLIDINGWINDOW:3:20 MINLEN:20). Cleaned reads were aligned to the *Drosophila melanogaster* reference genome r6.47, Flybase release FB2022_04 using the RNA-seq aligner, STAR version STAR_2.7.3a (Dobin et al., 2013). Read pairs mapping concordantly and uniquely to the exon regions of the annotated genes were counted using featureCounts tool ver. 2.0.0 ((Liao et al., 2014) of subread package. Read alignments to the antisense strand, or to multiple regions on the genome or those overlapping with multiple genes were ignored (parameters: -s 2 -p -B -C). Differential expression analysis was performed using DESeq2 ver. 1.24.0 (Love et al., 2014) and the p-values were corrected for multiple-testing using the Benjamini–Hochberg method. Raw reads and processed counts are available on NCBI GEO at accession GSE335328. All library preparation and sequencing were performed at the IU Center for Genomics and Bioinformatics (CGB).

### Sgs-GFP imaging

*w[*]; P{w[+mC]=Sgs3-GFP}3* (RRID:BDSC_5885) L1 larvae were placed on PFOA supplemented media for 84 h, after which time they were removed from the food, washed with cold 1x PBS, and placed on ice to immobilize. Larvae were subsequently imaged using a Leica MZ10F microscope with an ET GFP filter and a Leica DFC 450 digital camera.

### Developmental gene analysis

DEGs identified by RNA sequencing were evaluated for developmental regulation during larval stage 3 (L3) using the modENCODE developmental RNA-seq dataset. Genes were classified as developmentally regulated if their expression changed by more than twofold (increase or decrease) between the “larva L3 12 h” and “larva L3 puff stage 1–2” timepoints (Roy et al., 2010). Comparisons were performed step wise between low (or lower) and moderate, moderate and moderately high, moderately high and high, high and very high, and very high and extremely high. All genes were reexamined to ensure there was a 2-fold change in relative expression levels with the cut off set to 10 or lower. The resulting gene set was further stratified into “low expression” or “moderate+ expression” categories based on their maximum expression levels in the modENCODE temporal expression dataset at either L3 timepoint.

### FOXO nuclear localization

FOXO localization was examined as previously described (Birnbaum et al., 2019). Briefly, fat bodies from third-instar (L3) larvae were dissected 84 h after egg laying (AEL) in 1× phosphate-buffered saline (PBS; pH 7.4). Tissues were fixed in 4% paraformaldehyde and subsequently blocked for 1 h at room temperature in PBS containing 0.1% Triton X-100 (PBST) and 5% normal goat serum (NGS). Dissected fat bodies were then incubated with an anti-FOXO primary antibody (1:1000 dilution, a kind gift from Hua Bai, Birnbaum et al., 2019) in PBST for 16 h at 4 °C with gentle agitation. Samples were washed and incubated with donkey anti-rabbit Alexa Fluor 488 secondary antibody (Invitrogen, A21206). Images were acquired using a Leica SP8 confocal microscope and processed with ImageJ Fiji.

### Insulin gene comparison

Lists of significantly altered genes identified by RNA sequencing (RNA-seq; Table S3-S4) in comparisons between unexposed controls and larvae exposed to either 0.036 µM or 3.6 µM PFOA were intersected with curated insulin-signaling gene sets. Insulin-related gene sets were obtained from the Drosophila RNAi Screening Center’s Gene List Annotation for Drosophila (GLAD) database and the FlyBase insulin-like receptor signaling pathway annotation (Hu et al., 2011, Öztürk-Çolak et al., 2024).

### JC-1 measurements

JC-1 analysis was conducted following the protocol described previously (Mahmoudzadeh et al., 2024). Briefly, fat bodies from third-instar (L3) larvae were dissected 84 h post egg laying (HPEL) in Grace’s Insect Medium. Dissected tissues were washed in PBS containing 0.1% Triton X-100 (PT) and incubated in the dark with 5 µM JC-1 (5,5′,6,6′-tetrachloro-1,1′,3,3′-tetraethylbenzimidazolylcarbocyanine iodide) dye in PBS. Following incubation, fat bodies were washed in PT, mounted in 80% glycerol, and imaged live. Confocal images were acquired using a Leica SP8 confocal microscope with 488 nm and 568 nm laser excitation to detect JC-1 monomers and J-aggregates, respectively. Image analysis was performed using ImageJ. For each fat body, fluorescence intensity was measured in ten defined regions of interest in the 488 nm channel, and the same regions were quantified in the 568 nm channel. The ratio of 568 nm to 488 nm fluorescence was calculated for each region to estimate mitochondrial membrane potential. Statistical significance was determined using an unpaired t-test in GraphPad Prism version 10.4.1.

### Calorimetry

After an initial synchronized egg lay, 50 embryos were transferred to exposure chambers with either 0 or 500 µM PFOA at 25°C and 50% humidity. Larvae were then resynchronized at the L2-L3 molt. Biocalorimetry measurements were performed the following day using a previously described apparatus (Panda et al., 2025, Panda et al., 2024, Hur et al., 2020), modified for performing larvae measurements along with remote optimal viewing (Figure 3C). Larvae were suspended in Graces’ Insect Basal Medium (Corning 13-200-CV). Before the heat output measurement of the larvae, Graces’ Insect Basal Medium was filled in the sensing capillary channel of the biocalorimeter using a syringe pump (Harvard Apparatus, Pump 11 Pico Plus Elite, see Fig. 3C). Individual larvae were loaded into the capillary channel and positioned near the sensing thermistor with the help of a stopper (see Fig. S4A in the SI). The stopper is made of a multimode optical glass fiber (Thor Labs, UM22-300) that also acts as a light source for imaging the sample, together with the optical windows assembled on the outer shield and an inverted optical microscope (Zeiss Axiovert 200 with 10X objective), as shown in Fig. 3C.

During a calorimetric measurement, the buffer was continuously moved through the sensing capillary channel at a flow rate of 8-20 microliters/minute (µl/min) to supply nutrients and oxygen to the larvae. The heat output from individual larvae were recorded, along with remote viewing of the sensor region to ensure that the larvae stayed near the sensor. If the larvae swam away from the sensor (thermistor), the larvae were brought back to the sensor by remotely controlling the syringe pump to minimize the thermal perturbation of the calorimetric setup. Reference calorimetric measurements without the larvae were taken under identical conditions (at the same buffer flow rate). Figure S4B shows the dark field optical image of the sensor region without any larvae during a reference measurement and then with one larva when its heat output was being measured. The average heat output for each larva was determined by averaging the heat output of the stabilized readings over ∼15 minutes of measurement, excluding the readings when the larvae moved away from the sensor or when the readings were stabilizing.

During a measurement, the sensing thermistor detects temperature changes (Δ*T*) in its vicinity, caused due to the metabolic output of a larva. The metabolic heat output (*Q̇*) from a larva can be estimated from the measured temperature change via *Q̇* = *G*_*th*_ × Δ*T* (Sadat et al., 2013), where *G*_*th*_ is the thermal conductance of the sensor to the ambient. The larvae heat output measurements were performed with three flow rates, i.e., 8, 15, and 20 µl/min. *G*_*th*_ was calibrated for each of the three flow rates by dissipating known amounts of heating power at the sensing thermistor and measuring the resulting temperature change (see Panda et al., 2025 for detailed measurement technique of *G*_*th*_). *G*_*th*_ was determined to be ∼316.6, ∼380.1, and ∼399.2 µW/K at 8, 15, and 20 µl/min flow rates, respectively (Figure S4C).

### Environmental sensitivity assays

For thermal sensitivity assays, groups of 50 embryos were transferred to BDSC food containing increasing concentrations of PFOA and incubated at 29 °C and 50% relative humidity. The number of pupae per vial was scored at 96 h after egg laying (AEL), and larval mortality was assessed by counting dead larvae on the sides of the vials at 184 h AEL.

Food hydration experiments were conducted using BDSC summer food prepared in October and November 2025, following recognition of potential seasonal variation in assay outcomes. Embryos were placed on BDSC summer food containing PFOA and supplemented with either 1% water (v/v; standard summer recipe) or an additional 5% water (v/v; winter recipe). Cultures were incubated at 25 °C with 50% relative humidity, and larval development rate was quantified by counting the number of pupae at 120 h AEL.

### PFOA quantification

For quantification, L1 larvae were exposed to PFOA and allowed to pupate in vials before being transferred to clean BDSC food, where they were collected 5-days post-eclosion. Adults were separated by sex and weighed with an analytical balance before being extracted with water-methanol and chloroform. The fractions were dried down and resuspended in HPLC grade methanol.

PFOA concentrations were quantified using a previously described method (Zheng et al., 2023). Briefly, samples were analyzed using an ultra-performance liquid chromatograph coupled with a triple-quadrupole mass spectrometer (Agilent 1290 Infinity II UPLC – 6470 QQQ-MS) in the negative electrospray ionization (ESI-) mode was used to analyze PFOA. Chromatographic separation was achieved on an Acquity UPLC BEH C18 column (50 mm, 2.1 mm i.d., 1.7 μm thickness, Waters) at 40°C. Mobile phases consisted of 2 mM ammonium acetate in water (A) and 2 mM ammonium acetate in methanol (B). The gradient was 10% B for 0.5 min initially, ramped to 40% B at 1 min, and then increased to 100% B at 17.5 min. The instrument was equilibrated for 3.5 min after every run. The injection volume was 5 μL. The nebulizer, gas flow, gas temperature, capillary voltage, sheath gas temperature, and sheath gas flow were set to be 25 psi, 10 L/min, 300°C, 2800 V, 330°C, and 11 L/min, respectively. Data acquisition was operated under dynamic multiple reaction monitoring (dMRM) mode.

### Metabolomic analysis

Oregon R embryos were hatched in bottles of BDSC food containing either 0, 0.036, or 3.6 μM PFOA. Larval samples were collected at 84 h AEL. Adult samples were collected by transferring 20 newly eclosed males and females from individual bottles to fresh control food and aged for 5 days, at which time each culture was sorted by sex, males and females individually transferred into pre-tared 2 mL screwtop bead tubes, and the sample mass recorded using a Mettler Toledo XS105 Analytical Balance. Sample tubes were then drop frozen in liquid N2 and stored at −80 °C until extraction.

Samples were transferred to a −20 °C enzyme carrier caddy and 0.8 mL of prechilled (−20 °C) 90% methanol containing 2 µg/mL succinic-d4 acid (Sigma-Aldrich; 293075) was added into each tube. Samples were homogenized at 4 °C using an Omni Bead Ruptor 24 homogenizer (30 seconds at 6.45 m/second). The homogenized samples were incubated at −20 °C for 2 hours and then centrifuged at 20000 x g for 5 min at 4 °C. 0.6 mL of supernatant was transferred to a new 1.5 mL microcentrifuge tube, taking care not to disturb the precipitant. 200 µL of the supernatant was subsequently pipetted to a well of a 96 well plate (SureSTART 60180-P217B). The sample plate was dried under a nitrogen air stream, sealed with a foil cover (VWR 60941-126), and stored at −80 °C until shipping.

UHPLC-MS metabolomics analyses were performed at the University of Colorado Anschutz Medical Campus, as previously described (Nemkov et al., 2019). Briefly, the analytical platform employs a Vanquish UHPLC system (Thermo Fisher Scientific, San Jose, CA, USA) coupled online to a Q Exactive mass spectrometer (Thermo Fisher Scientific, San Jose, CA, USA). The (semi)polar extracts were resolved over a Kinetex C18 column, 2.1 x 150 mm, 1.7 µm particle size (Phenomenex, Torrance, CA, USA) equipped with a guard column (SecurityGuard^TM^ Ultracartridge – UHPLC C18 for 2.1 mm ID Columns – AJO-8782 – Phenomenex, Torrance, CA, USA) using an aqueous phase (A) of water and 0.1% formic acid and a mobile phase (B) of acetonitrile and 0.1% formic acid for positive ion polarity mode, and an aqueous phase (A) of water:acetonitrile (95:5) with 1 mM ammonium acetate and a mobile phase (B) of acetonitrile:water (95:5) with 1 mM ammonium acetate for negative ion polarity mode. The Q Exactive mass spectrometer (Thermo Fisher Scientific, San Jose, CA, USA) was operated independently in positive or negative ion mode, scanning in Full MS mode (2 μscans) from 60 to 900 m/z at 70,000 resolution, with 4 kV spray voltage, 45 sheath gas, 15 auxiliary gas. Calibration was performed prior to analysis using the Pierce^TM^ Positive and Negative Ion Calibration Solutions (Thermo Fisher Scientific).

### Metabolomic data analysis

Data were normalized to sample mass and the succinic-d4 acid control prior to analysis. Any compound that was undetectable in 2 or more replicates (value of 0) for any of the three exposure conditions were removed, the exception being if a metabolite was not detected in 5 or more of the 6 replicates for one condition but was detectable in at least 5 replicates of the other conditions. The normalized and filtered data was then analyzed using MetaboAnalyst 6.0. Data underwent log transformation and Pareto scaling prior to analysis. Heatmaps in Figure 6 were produced in GraphPad Prism 10 and metabolites rank ordered according to F-value.

### Targeted acyl-carnitines analysis

Targeted measurements were performed for two acyl-carnitines for which authentic standards were available: palmitoyl-L-carnitine (C16:0; Avanti Polar Lipids, 870323P) and oleoyl-L-carnitine (C18:1; Avanti Polar Lipids, 870321P). Larval PFOA exposures, adult collection, and sample handling were performed exactly as described for the semi-targeted metabolomic analysis above.

Lipids were extracted using a biphasic organic solvent extraction. Samples were placed in a pre-chilled enzyme caddy, and 400 µL of methanol containing internal standards in PBS (225:188, v/v) was added to each sample, along with a blank. Samples were homogenized at 4 °C using an Omni Bead Ruptor 24 homogenizer for 30 s at 6.45 m s⁻¹. Homogenates were transferred to 1.8 mL glass screw-cap vials containing 750 µL of HPLC-grade methyl tert-butyl ether (MTBE) and incubated on ice for 1 h, vortexing every 15 min, followed by centrifugation at 3220 rcf for 10 min at 4 °C. The organic phase was transferred to a 1.5 mL Eppendorf tube. The remaining aqueous phase was re-extracted with an additional 750 µL of MTBE/MeOH/Milli-Q H₂O (10:3:2.5, v:v:v), vortexed, incubated on ice for 10 min, centrifuged at 3220 rcf, and the organic phase was combined with the initial extract. Combined organic extracts were dried under a nitrogen airstream and stored at −80 °C.

Dried samples were reconstituted in 200 µL of water (Fisher Optima W6500), vortexed, diluted with 800 µL of acetonitrile (Fisher Optima A955-500), and vortexed again. Samples were centrifuged at 15,000 rcf for 15 min at 4 °C. From each sample, 200 µL of supernatant was used for analysis, and 20 µL was reserved to generate a pooled quality-control (QC) sample.

Samples (2 µL injections) were analyzed using an Agilent 1290 Infinity II LC system (Multisampler G7167B, Column Compartment G7116B, and Binary Pump G7120A) coupled to an Agilent 6495C triple quadrupole mass spectrometer. Chromatographic separation was performed on an Agilent Poroshell 120 HILIC-Z column (2.1 × 150 mm, 2.7 µm). Solvent A consisted of 20 mM ammonium acetate/ammonium hydroxide buffer with 5 µM medronic acid (Agilent InfinityLab Deactivator Additive), pH 9.2, and solvent B was 100% acetonitrile. The column compartment was maintained at 15 °C throughout the analysis. An initial flow rate of 0.4 mL/min of 90% B was set for one min and ramped to 0.4 mL/min of 83% B over the course of 7 mins. This was held for 1 minute before being ramped to 0.4 mL/min of 10% B over the course of 0.1 mins. The flow of 0.4 mL/min of 10% B was held constant for 3 mins before being ramped to back to 0.4 mL/min of 90% B over the course of 1 min with the flow rate being increased to 0.5 mL/min and 90% B over the course of 0.1 mins and kept this flow rate and ratio steady for 2.8 mins.

The LC gradient and MS acquisition parameters were as described above. The mass spectrometer was operated in positive-ion dynamic multiple reaction monitoring (dMRM) mode. Analytes included oleoyl-L-carnitine (C18:1) and palmitoyl-L-carnitine (C16:0), with d9-labeled carnitine analogs used as internal standards. Parent ions corresponded to intact molecular ions (no protonation). Quantification was based on the most intense transition for each compound and verified using a qualifier ion. Ion counts were normalized to the mass of extracted fly samples.

### Statistical Analysis

Data analysis was conducted using GraphPad Prism v11.0. For each dataset, normality was assessed using the Shapiro–Wilk test. For datasets that met assumptions of normality and homogeneity of variance, statistical significance was determined using parametric tests (Student’s t-test for two-group comparisons or one-way/two-way ANOVA for multi-group comparisons) followed by appropriate post hoc tests as indicated in the figure legends. When data did not meet these assumptions, a Kruskal–Wallis test with multiple comparisons was used.

## ACKNOWLEDGEMENTS

We thank the Bloomington Drosophila Stock Center (NIH P40OD018537) for providing fly stocks, Flybase (NIH 5U41HG000739), the Indiana University Light Microscopy Imaging Center, the Indiana University Center for Genomics and Bioinformatics, the Indiana University Mass Spectrometry Laboratory, Dr. Hua Bai for providing us with the FOXO antibody, Karen Yannell of Agilent Technologies for the base LC method from which the chromatography to measure acyl-carnitines was adapted, and Shane Tichy of Agilent Technologies for the generous donation of the 6495C QQQ instrument through Agilent’s partnership in the Center for Bioanalytical Metrology. The biocalorimetry experiments were supported by National Institutes of Health NIGMS R35GM133737 grant to S.Y., the Alfred P. Sloan fellowship, the McKnight scholar grant, and the Chan Zuckerberg collaborative pairs grant to S.Y. The JC1 and biocalorimetry studies were supported by the National Institute of General Medical Sciences of the National Institutes of Health under a R35 Maximizing Investigators’ Research Award (1R35GM119557) to J.M.T. Metabolomic studies were supported by the National Institute Of Diabetes And Digestive And Kidney Diseases of the National Institutes of Health under Award Number R01DK136945 to T.N., A.D., and J.M.T. The Drosophila studies were partially supported by the Bloomington Drosophila Stock Center, which is funded by National Institutes of Health grant P40 OD018537. This project received funding from the European Union’s Horizon 2020 Research and Innovation program under Grant Agreement No. 965406. The work presented in this publication was performed as part of the ASPIS Cluster. This output reflects only the authors’ views, and the European Union cannot be held responsible for any use that may be made of the information contained therein. This publication was also made possible with support from the Indiana Clinical and Translational Sciences Institute, which is funded in part by Award Number UL1TR002529 from the National Institutes of Health, National Center for Advancing Translational Sciences, Clinical and Translational Sciences Award. The content is solely the responsibility of the authors and does not necessarily represent the official views of the National Institutes of Health.

