## Supplemental Figures 1-4 for "Exposure to perfluorooctanoic acid accelerates *Drosophila melanogaster* juvenile development and disrupts mitochondrial metabolism"

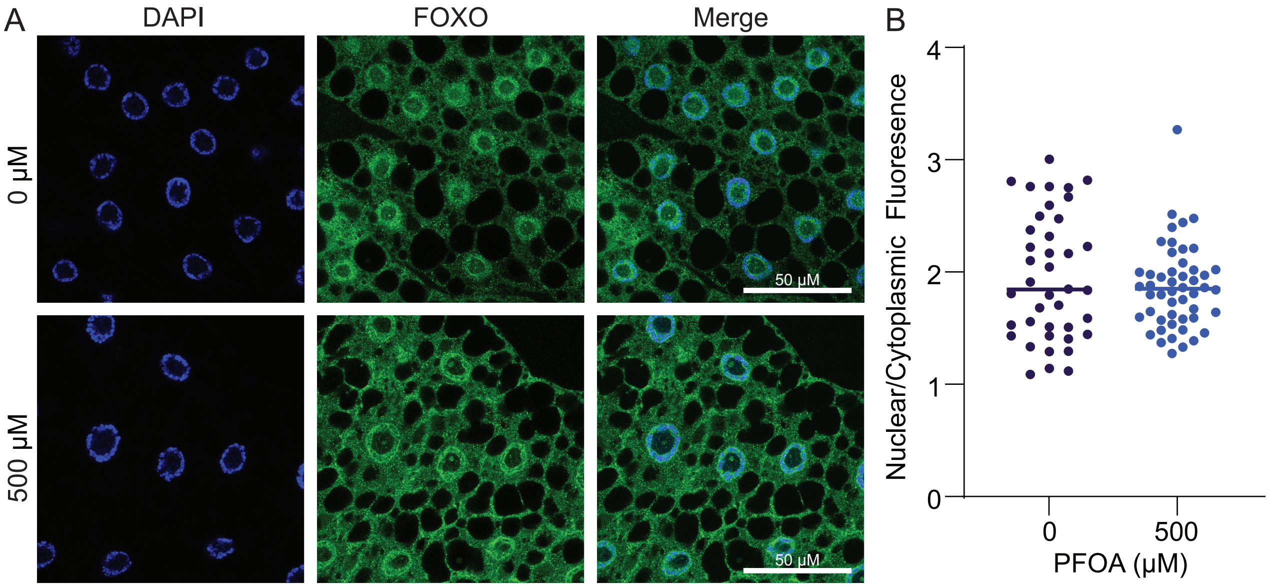


**Figure S1. FOXO subcellular localization following PFOA exposure.** Fat bodies were dissected from L2 larvae at 60 h after egg laying (AEL). Tissues were immunostained using an anti-FOXO antibody. (A) Representative images of fat bodies from larvae exposed to 0 µM or 500 µM PFOA, showing DAPI and FOXO channels individually and merged. (B) Quantification of the nuclear-to-cytoplasmic FOXO ratio measured using ImageJ. Each data point represents a field of view from 4-5 fat bodies. Data were analyzed using a Kruskal–Wallis test with multiple comparisons. No significant differences were found.

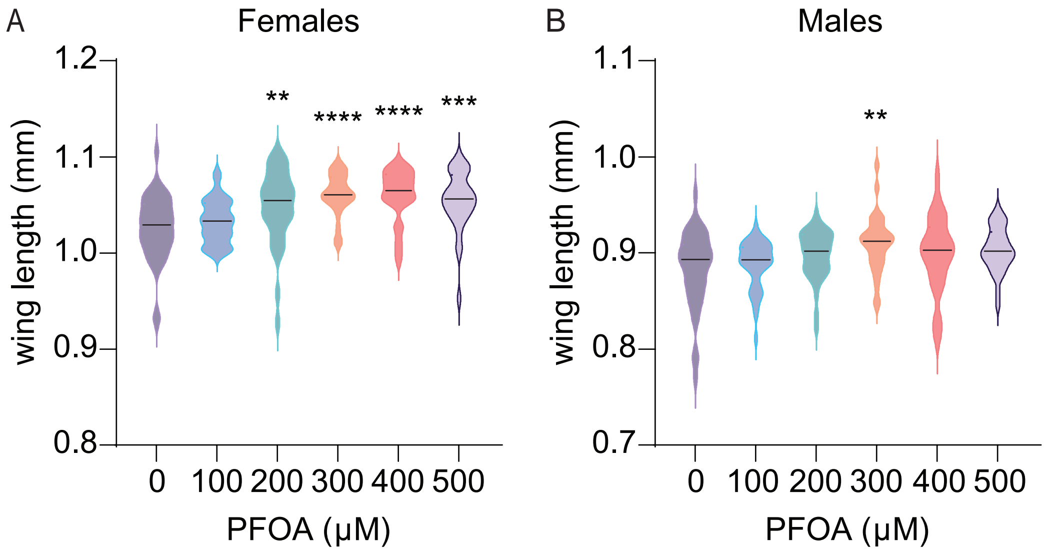


**Figure S2. Effects of PFOA exposure on adult wing size under elevated temperature conditions.** Oregon R larvae were reared at 29 °C and exposed to increasing concentrations of PFOA. Wing lengths were measured in adult (A) females and (B) males 1 day after eclosion (n=24-40 wings). Data were analyzed using Kruskal–Wallis tests with multiple comparisons to the 0 µM control. *$p < 0.05$. **$p < 0.01$. ***$p < 0.001$. **** $p < 0.0001$.

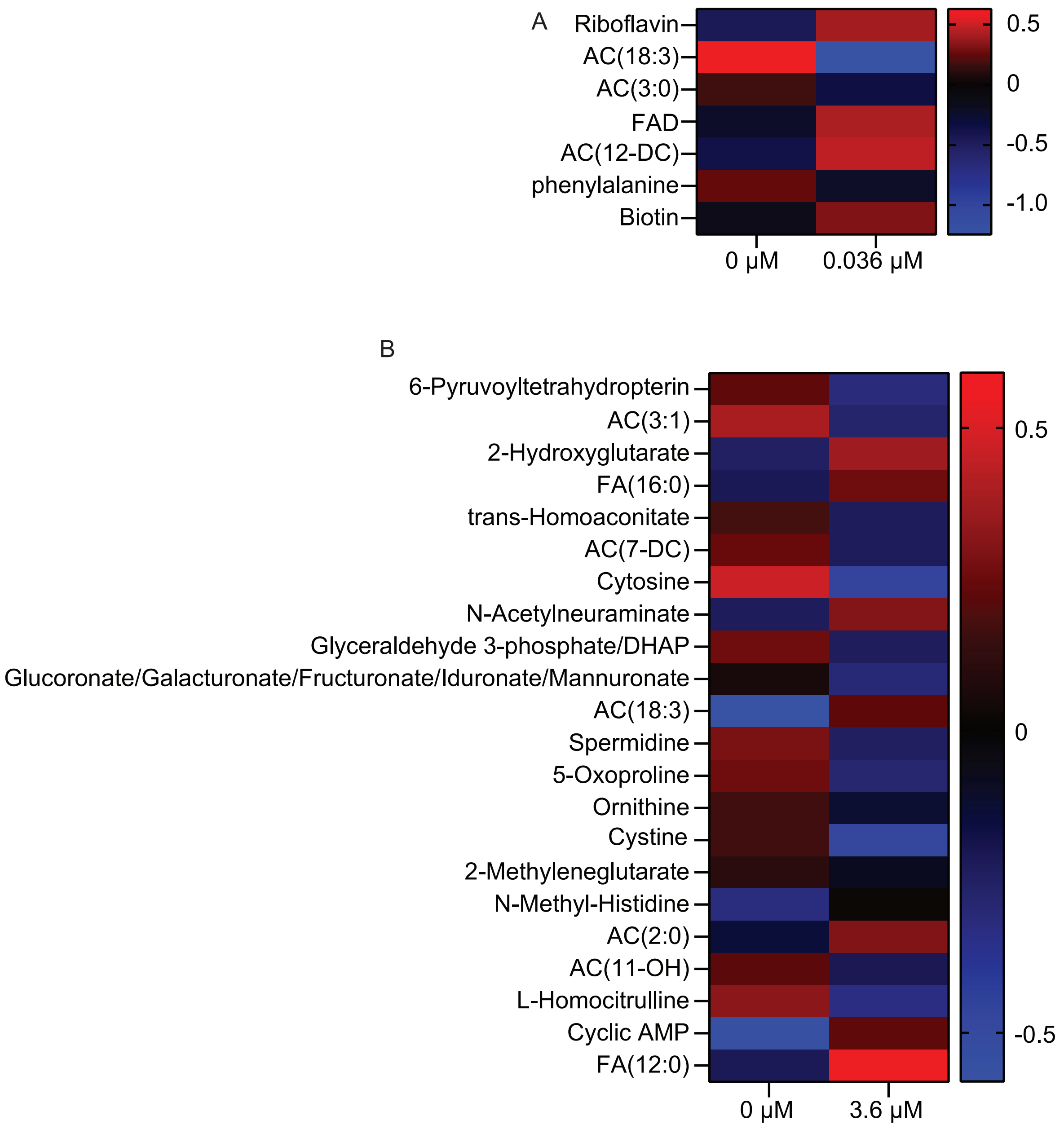


**Figure S3. PFOA exposure has limited effects on the larval metabolome.** Semi-targeted metabolomics was performed on whole larvae exposed to 0, 0.036 μM and 3.6 μM PFOA. (A,B) Heatmaps showing the metabolites that exhibited a statistically significant change $p < 0.05$ following developmental PFOA exposure. Data was normalized to sample mass and the spike-in internal standard. Data analysis conducted with Metaboanalyst 6.0. Metabolites were measured by LC–MS at 84 h AEL using 5 biological replicates per condition, each consisting of 25 larvae. Statistical analyses were conducted using MetaboAnalyst 6.0.

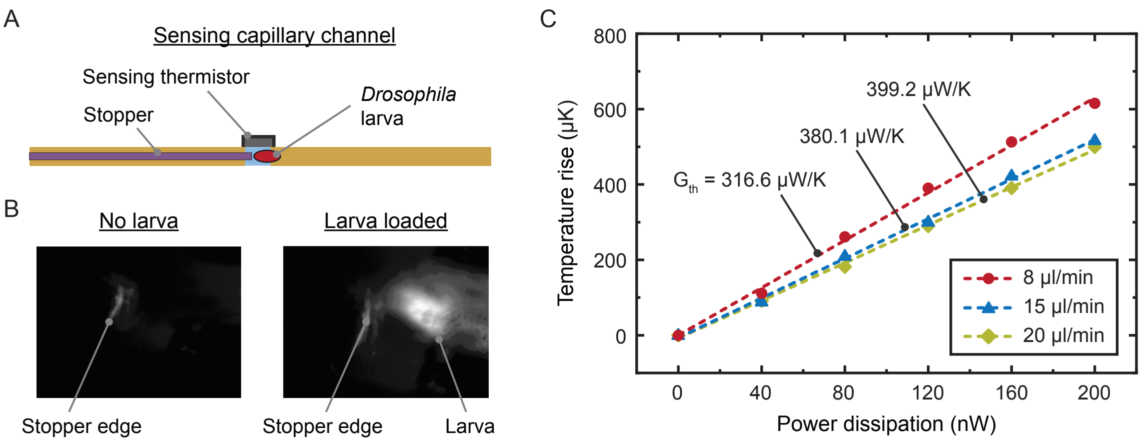


**Figure S4. Biocalorimetry thermal conductance measurements.** (A) Schematic of the sensing capillary channel of the biocalorimeter, indicating the stopper, sensing thermistor, and the position of the larvae during heat output measurements. (B) Dark-field optical microscopy images of the sensor region without a larva and with a single larva positioned near the stopper. (C) Thermal conductance (G_th) of the sensing capillary at different medium flow rates, measured to be approximately 316.6, 380.1, and 399.2 µW/K at flow rates of 8, 15, and 20 µL/min, respectively.
